# Morality recruits neural reward circuitry to shape economic decision making

**DOI:** 10.1101/2020.01.27.920694

**Authors:** Jie Liu, Xiaoxuan Huang, Chong Liao, Fang Cui

## Abstract

The present study combined a novel hypothetical investment game with functional magnetic resonance imaging systemtically examined how morality modulates economic decision making in decision phase and outcome phase. We manipulated the morality of the investments by choosing each investment project based on subjective ratings on their moral valence and social benefits. There were three categories of investment morality: Green (moral), Red (immoral), and Neutral. The behavioral and neural responses during the investment decision and outcome phases were recorded and compared. Results showed that: behaviorally, people are willing to invest a larger amount of money into a moral project that may benefit society than they are into an immoral project that they think will harm society. They also rate gains in moral investments as more pleasant and losses as the most unpleasant. In the brain, we found that the reward system, especially the bilateral striatum, was involved in modulating functional connectivity during both phases, but in different ways. During decision making, the functional connectivity between fusiform gyrus and striatum might underlie the observed investing bias (Green over Red projects), while the covariation of BOLD signals in bilateral striatum with the behavioral tendency might explain the effect observed during the outcome evaluations. Our study provides evidence that morality modulates both the decision making and the outcome evaluation in economic situations.

## Introduction

A growing literature from psychology, sociology, and neuroscience suggests that morality is deeply embedded in our daily decision making. Even in the realm of economics, people are not solely motivated by self-interest—as proposed by rational decision theorists—but are also motivated by moral considerations (Lewis and Mackenzie, 2000). For instance, Responsible Investments (RIs), which refer to investments aiming to maximize profit while taking moral concerns into account, have grown significantly all over the world (Derwall et al., 2011; Riedl and Smeets, 2017). This indicates morality’s influence in traditionally “pure” profit-seeking activities.

Many social dilemmas that have been investigated deeply in the fields of social psychology and social neuroscience involve conflicts between moral values and economic interests. For example, the Dictator Game (DG) examines the conflict between “being fair” and “giving more money to oneself” and Third-party Punishment (TPP) looks at “punishing the bad guy” vs. “keeping more money for oneself”. Imaging studies have found that neural activity in regions related to reward processing, including the striatum, ventromedial prefrontal cortex (vmPFC), amygdala, and orbitofrontal cortex (OFC) was correlated with cooperative tendency in such social behavior (for a review see Strang and Park, 2017). In both the DG and TPP, bilateral striatum was activated when individuals made prosocial choices (i.e., allocated more money to the receivers in the DG or altruistically punished the free-riders in the TPP) (Cornelissen et al., 2011; Strobel et al., 2011; Wei et al., 2017). A recent study created a conflict between a common immoral behavior (“Harming others”) and participant monetary benefits and found that participants with stronger moral preferences in this conflict had lower activity in the dorsal striatum in response to money gained from harming others (Crockett et al., 2017). Another recent study exploring how the brain solves the conflict between moral value and monetary value reported that the insula and lateral prefrontal cortex engage in weighing monetary benefits and moral costs while the putamen engages in weighing monetary costs against compliance with one’s moral values. Thus, the authors suggested that reward-related regions, especially the striatum, are involved in processing both moral and economic values, and that these representations interact with each other (Qu et al., 2019). These findings indicate that reward circuitry might play a significant role in resolving the conflict between morality and economic interests.

Conflicts between morality and money do not always exist. Most of the time, morality is simply one of several motivational concerns that influence our daily decisions and does not necessarily conflict with monetary interests. One example of this situation is the devaluation effect of moral contagions. Previous studies have found that “moral contagions” affect economic decisions related to the “contaminated” objects such that individuals value objects more if they are associated with respectable people and less if they are associated with immoral people (Hood et al., 2011; Kramer and lock, 2011; Newman and Bloom, 2014). An fMRI study reported that the devaluation effect was supported by activation of the putamen and its connectivity with the vmPFC (Liu et al., 2019). Another study found that in a trust game, prior moral information about trading partners modulated activation in the dorsal striatum during both the decision phase and the outcome phase (Delgado et al., 2005). These results suggest that reward circuitry underlies the modulating effect of morality in these types of economic decisions on a broader level.

The hypothesis that morality shapes economic decision making by recruiting reward circuitry is consistent with the “common currency” theory (Levy and Glimcher, 2012; Sescousse et al., 2015), which assumes that a single neural circuit determines the motivational significance of both monetary and non-monetary events. The activity of this circuit represents the integrated value of all factors that are relevant for a choice. Thus the “moral value” and “monetary value” are represented on a common currency scale and are interpreted as a subjective value for choices (Caspers et al., 2011; Cushman, 2013; Sescousse et al., 2015). Evidence has been found to support this hypothesis. For instance, studies have found that striatal neurons are sensitive to the expected “moral value” in moral judgment tasks (Decety and Porges, 2011; Shenhav and Greene, 2010; Yoder and Decety, 2014). Additionally, the striatum was found to be activated when giving charitable donations (Harbaugh et al., 2007), performing altruistic punishment (de Quervain et al., 2004), and making other types of moral decisions (Fumagalli and Priori, 2012).

Although several studies have alluded to the importance of the reward system in allowing morality to modulate economic decision making, few studies have systematically tested this hypothesis. Besides, reward circuitry has been suggested to be involved not only in representing expected subjective value during the decision phase but also in representing experienced subjective values during the outcome phase (Bartra et al., 2013). How morality modulates economic decision making in both phases remains unclear.

In the present study, participants were asked to play a hypothetical investment game in which they were presented with a project’s information, including the investment category and the project code. The probability to either gain or lose 5% of the invested money was different for each project. Participants decided how much they would like to invest. We manipulated the morality of the investments by choosing each project based on subjective ratings they had received earlier on their moral valence, social benefits, and social reputation. There were three categories of investment morality: Green (moral), Red (immoral), and Neutral. Gain probability was balanced across the moral directions. Participants were encouraged to make as much profit as possible because their actual payment for participating in the experiment would be calculated based on their performance in the game. In this way, we were able to examine how morality shapes economic decision making in a relatively implicit matter. The behavioral and neural responses during the decision and outcome phases were recorded and compared among the different moral valences (Green vs. Red vs. Neutral).

We hypothesized that: 1) during the decision phase, participants would show a preference to invest in Green projects rather than Red projects. We predicted that this behavioral bias would correlate with neuronal activity and connectivity within regions of the reward network, such as the striatum, reflecting the effect of morality in calculating subjective value during decision making; 2) during the outcome phase, morality would modulate the subjective ratings of pleasantness as well as the neuronal responses in reward-related regions when observing the investment outcome, thus reflecting the effect of morality on feedback evaluation.

## Methods

### Ethics

The study was conducted according to the ethical guidelines and principles of the Declaration of Helsinki and was approved by the Medical Ethical Committee of Shenzhen University Medical School. Informed consent was obtained from all participants after they fully understood the procedures. Participants were remunerated for their time (about 7 USD/hour) and received their game earnings.

### Participants

Thirty-five right-handed participants recruited from Shenzhen University participated in the fMRI experiment. Participants were screened for a history of neurological disorders, brain injury, and developmental disabilities. All had normal or corrected-to-normal vision. Two participants who had excessive head movements (> 2° in rotation or >2 mm in translation) during the scanning were excluded, leaving 33 participants for data analysis (15 females; age: 22.2 ± 3.7 y).

### Pretest questionnaire

In order to determine the investment categories for the fMRI experiment, we designed a pretest questionnaire and sent it to a list of 50 participants (healthy adults from Shenzhen University; 25 females; age: 23.2 ± 1.6 y, who did not participate in the fMRI experiment). This list included 18 investment categories. Participants were instructed to rate their willingness to invest (*willingness to invest*), how familiar they were with the investment category (*familiarity*), how much they thought the investment would benefit society (*social benefits*), how moral they thought this kind of investment was (*morality*), how likeable the investors were who focused on that category of investments (*investor likeabilit*y), and how good the profit outlook for investment seemed (*profit outlook*) on 7-point Likert-like scales.

Correlation analysis revealed that the ratings for *social benefits*, *morality*, and *investor likability* were positively correlated with those for *willingness to invest* (*r* = 0.622, *p* < 0.001; *r* = 0.510, *p* < 0.001; *r* = 0.603, *p* < 0.001, respectively). These results suggested that behaviorally, moral concerns do modulate the willingness to invest such that investments that are more moral, more beneficial to society, and that have more morally reputable investors promote personal willingness to invest.

Nine investment categories were selected for the fMRI experiment based on the pretest ratings. First, categories with average familiarity scores ≥ 3 were discarded to exclude unfamiliar investment categories. Then, looking at the *moral* category, the three items with the highest ratings were selected as Green investments, the three with the lowest ratings were selected as Red investments, and the three in the middle were selected as Neutral investments. Thus, the Green investment categories were *Sewage Purification*, *Manufacture of Materials to Protect the Environment*, and *Green Food*, Red investment categories were *Gambling Industry*, *Tobacco Industry*, and *Real Estate*, and Neutral categories were *Clothing Industry*, *IT Indust*ry, and *Construction Businesses* (Fig.1A). One-way ANOVAs showed that ratings differed significantly across the three moral categories (Red, Green, Neutral) for *morality*, *social benefits*, and *investor likability* (*ps* < 0.001), but not for *familiarity* or *profit outlook* (*ps* > 0.352).

**Figure 1.**
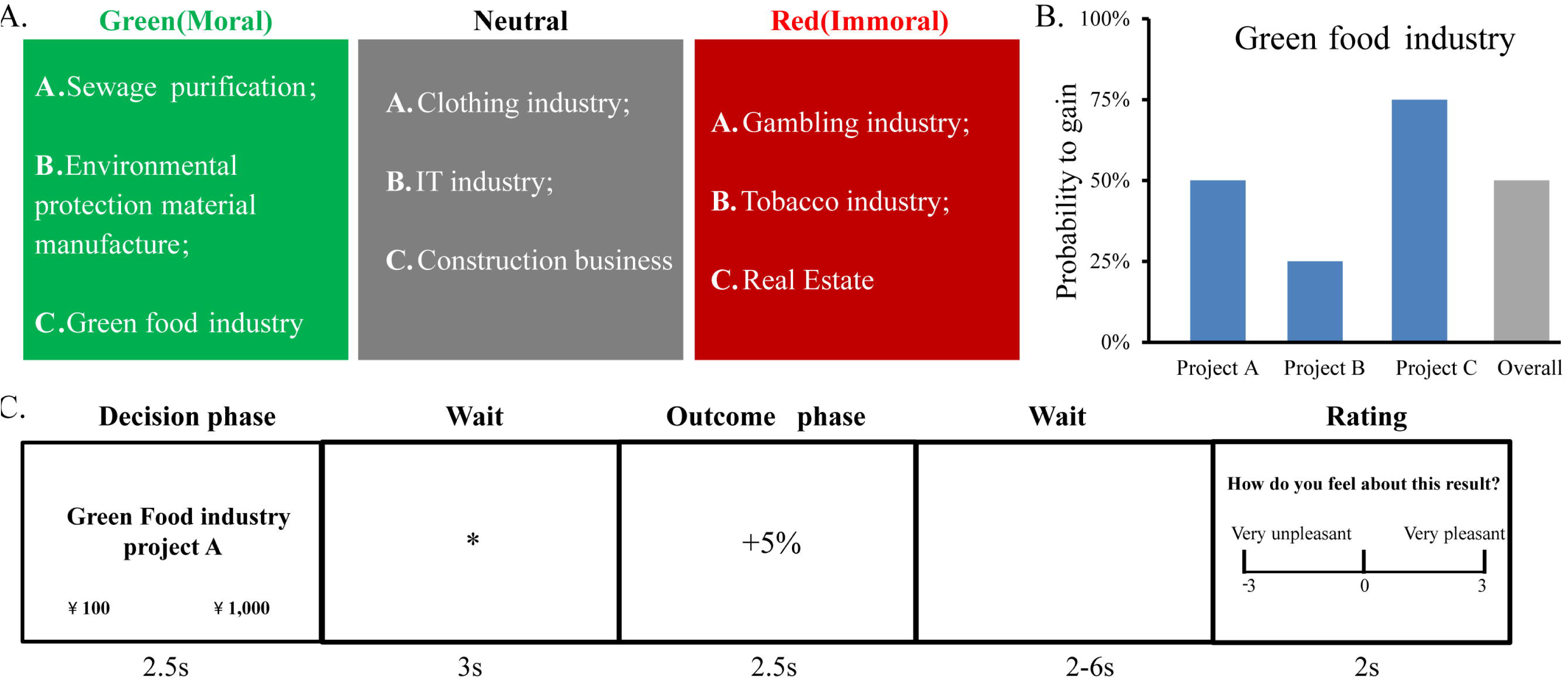
Experimental design. **(A)**. Moral valence of the investment categories: Green, Red, or Neutral based on data from a pretest. **(B)**. Example of the winning probabilities for each of the three projects within a category. In all cases, 25%, 50% and 75% winning probabilities were assigned pseudorandomly to the three projects. The overall winning probability of each category was always 50%. **(C)**. Trial structure. Participants had 2.5 s to decide to invest ¥100 or ¥1,000 into the presented project. After a 3 s wait, the outcome of investment (gain or loss) was presented for 2.5 s. After a 2–6 s wait, they were instructed to rate their feelings about the outcome.

### Experimental design and procedures

Participants were informed that they would play a hypothetical investment game. They were encouraged to make more money in the game. The more they earned in the game, the more they would be paid extra at the end of the experiment.

The fMRI experiment had a 3 (moral valence of the investment category: Green, Red, or Neutral) × 2 (Investment amounts: 1,000 RMB or 100 RMB) within-subject design. Participants were shown information about each investment project including the category to which it belonged and an alphabetic serial number (A, B, or C) indicating different projects. They faced two options: to invest a large amount (¥1,000) or a small amount (¥100) into the given project. They were told that they would either gain or lose 5% of the invested money.

Each category had three projects (e.g., *Green food* project A, B, and C), one of which had a 25% chance of gain, one a 50% chance, and the other a 75% chance. All categories had an overall gain probability of 50%. Thus, the Green, Red, and Neutral investments had equal chances to make a profit (Fig. 1B).

Each trial began by presenting the project information for 2.5 s during which participants were required to indicate their decisions (i.e., to invest ¥1,000 or ¥100). The locations of the two options were randomized left or right trial by trial. After a 3-s fixation, participants were presented with the outcome (i.e., whether they gained or lost 5% of the invested money), which lasted 2.5 s. After a 2–6 s waiting period, in 25% of all trials, participants were instructed to rate how pleasant they felt on a 7-point scale (from very unpleasant [−3] to very pleasant [3]) (Fig.1B).

To make their choices, participants pressed the left or right buttons of an MRI-compatible button box. Each project was repeated 8 times, resulting in a total of 216 trials (9 categories × 3 projects × 8 repetitions). The experiment was divided into 4 runs of 54 trials and lasted approximately 1 h. Trials in each run were pseudorandomized such that the numbers of Green, Neutral, and Red investment categories were the same (18 trials/category × 4 runs = 72 trials/category). Before the scanning, participants were familiarized with the task using a practice block consisting of 12 trials.

### fMRI data acquisition and preprocessing

We used a Siemens TrioTim 3.0T MRI machine for data acquisition. Functional volumes were acquired using multiple slice T2-weighted echo-planar imaging (EPI) sequences with the following parameters: repetition time = 2,000 ms; echo time = 25 ms; flip angle = 90°; field of view = 192 mm; 41 slices covering the entire brain, slice thickness = 3 mm.

fMRI data were preprocessed in SPM12 (Wellcome Department of Imaging Neurosciences, University College London, U.K., http://www.fil.ion.ucl.ac.uk/spm). Images were slice-time corrected, motion-corrected, and normalized to Montreal Neurological Institute (MNI) space for each individual with a spatial resolution of 3 mm × 3 mm × 3 mm. Images were then smoothed using an isotropic 4-mm Gaussian kernel and high-pass filtered at a cutoff of 128 s. A threshold of *p* < 0.001, uncorrected with a voxel size > 10 was considered statistically significant in the whole-brain analyses. Additionally, we used cluster-based *FWE* correction for multiple comparisons. Significance was set to the corrected *p* < 0.05.

### fMRI data analysis

#### Decision phase

##### GLM

Statistical parametric maps were generated on a voxel-by-voxel basis with a hemodynamic model to estimate the brain response. Presentation of project information, the interstimulus intervals, the feedback of the investment, the rating (if any), as well as inter stimulus interval (ISI), were included in the general linear model (GLM) at the single-participant level. The six rigid-body parameters were also included in the GLM to exclude head-motion nuisance. We generated two general linear models to explore how morality influences economic decisions. GLM1 was set up to localize the brain regions that were correlated with economic decisions. We considered the trials in which the participant invested ¥1,000 or ¥100 as independent regressors and defined the investment decisions as the contrast: ¥1,000 > ¥100. GLM2 was set up to explore brain regions that were sensitive to the moral valence of the project when making economic decisions. The independent regressors in GLM2 were the three moral valences of the investments (Red, Neutral, and Green). For the group-level analysis, a one-sample t-test was conducted using the whole brain as the volume of interest.

##### Localization of the Regions of Interest (ROIs)

We first localized several brain regions related to the investment decisions in GLM1 and considered the peaks of the significantly activated clusters as regions of interest. Spheres (radius = 6 mm) that were centered at these peaks were drawn. For each participant, we then calculated mean parameter-estimates of the three different moral valences of the investment categories in GLM2 within the spheres. We then performed one-way ANOVAs on the extracted values to determine the brain regions that were sensitive to moral valence.

##### Psychophysiological interaction (PPI) analysis

We used PPI analysis to explore the functional connectivity between ROIs that were sensitive to moral valence among brain circuits that were correlated with the behavioral decisions and the whole brain. For each participant, the mean time series for the seed region was extracted and adjusted using the *F*-contrast for the regressors. The first eigenvariate for the single-participant time courses was extracted from the seed region, using a sphere with a 6-mm radius that was centered at individual maxima and adjusted for the effects of interest. The GLM included the following PPI regressors: (1) the main effect of seed-region activity, (2) the main effect of the contrast between Green and Red directions, and (3) their interaction. These regressors corresponded to PPI.Y, PPI.P, and PPI.ppi in the design matrix. The six head-motion parameters were also entered into the GLM as covariates to regress out any head-motion artifacts. Low-frequency drifts in signal were removed using a high-pass filter with a cutoff at 128 s. After model estimation, the PPI.ppi regressor for Green minus Red was entered into a random-effects analysis for the entire group.

##### Correlation between functional connectivity (FC) and the behavioral index

Because we observed at the behavioral level that participants tended to invest more money in Green categories than in Red categories, we used the difference in the percent of ¥1,000 trials between Green and Red categories (i.e., Green_percent¥1,000_ – Red_percent¥1,000_) as the *behavioral index* to indicate how much decisions were modulated by morality.

By correlating functional connectivity strength with the behavior index, we were able to explore how brain functional connectivity supports the process in which morality influences economic decisions. We downloaded the results of a reward-processing meta-analysis from Neurosyth (https://www.neurosynth.org/) and used it as a mask (t-value ≥5). We selected the peaks of the significant PPI results among the mask as ROIs and conducted a correlation analysis between FC and the behavioral index.

#### Outcome phase

### GLM

We set up GLM3 to explore how morality interfaces with the outcome of the economic decision. Specifically, we investigated how brain activity that supports morality influenced outcome processing for Green and Red investment categories. Gain and Loss outcomes were considered as independent regressors in GLM3 at the single-participant level. In the second-level analysis, we considered the behavioral index as a regressor and regressed it against the contrast between Gain and Loss. Using the same meta-analysis mask, we selected the peaks of the significant brain activation results among the mask as ROIs and conducted a correlation analysis between the brain activity and the behavioral index.

## Results

### Behavioral results

#### Decision phase

##### Choice preference

To investigate how the moral valence of the investment category modulated investment behavior in a hypothetical investment game, we performed a one-way ANOVA on the percentages of participants who chose to invest ¥1,000 rather than ¥100 in all trials across three conditions (Green, Red, and Neutral). Results showed that the moral valence significantly modulated investment decisions (*F*_(2, 64)_ = 18.163, *p* < 0.001, η_p_^2^ = 0.36) such that Green projects received the most investments and Red projects received the least (Green: 0.65 ± 0.05, Neutral: 0.52 ± 0.05, Red: 0.34 ± 0.04; Green vs. Neutral: *p* = 0.008; Red vs. Neutral: *p* < 0.001; Green vs. Red: *p* < 0.001, Bofferroni corrected). No significant effect was observed for reaction time (*ps* > 0.184) (Fig. 2A).

**Figure 2.**
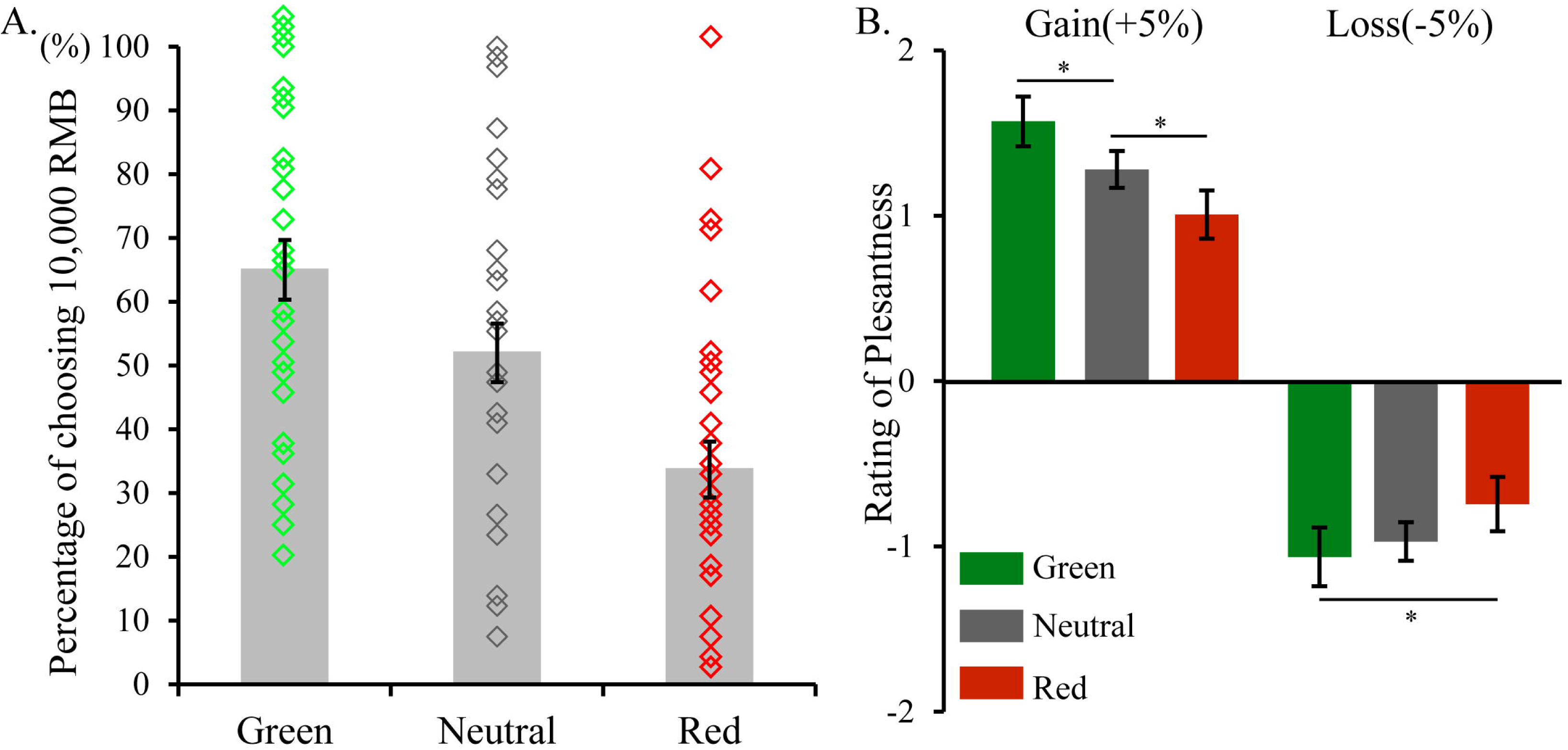
Behavioral results. **(A)** The results of the decision phase for each type of investment. **(B)** Pleasantness outcomes plotted by loss and gain for each type of investment (\**p* < 0.05).

##### Task Performance

Although the present study did not focus on feedback-learning, we nevertheless calculated the performance (i.e., how much money they gained in the game) for each moral category of investment. One-way ANOVA revealed significant differences between Green, Red, and Neutral conditions in performance (*F*_(2, 96)_ = 12.82, *p* < 0.001). Participants made the most profit for Green categories and the lease profit in Red categories (Green vs. Neutral: *p* = 0.005, Green vs. Red: *p* < 0.001, Red vs. Neutral: *p* = 0.034).

#### Outcome phase

We calculated ratings outcomes for each moral category and each outcome separately. Thus, six levels were generated: *Green_gain*, *Green_loss*, *Red_gain*, *Red_loss*, *Neutral_gain*, and *Neutral_loss*. We note that there was an imbalance in choosing ¥1,000 among Green, Red, and Neutral projects such that participants chose ¥1,000 more for Green projects. Mathematically, this means that the gain/loss for Green projects was higher than that for Red projects. To solve this problem, we used the percentage of each option closed across all trials as the weights in calculating subjective ratings. For example, the rating for Green_gain was calculated as (*Gain for Green choosing ¥1,000 × Percentage of choosing ¥1,000 in all Green trials*) *+* (*Gain for Green choosing ¥100 × Percentage of choosing ¥100 in all Green trials*).

Then we performed 3 (Moral valence: Green, Red, Neutral) × 2 (outcome valence: gain vs. loss) repeated-measure ANOVA on the weighted pleasantness ratings. We found a significant main effect of outcome (*F*_(1, 32)_ = 104.078, *p* < 0.001, η*_p_* = 0.765) such that participant felt better when they gained money than when they lost. Although we did not observe a significant main effect of moral valence (*p* = 0.273), the moral valence × outcome interaction was significant (*F*_(2, 64)_ = 9.171, *p* < 0.001, η*_p_* = 0.223). Pairwise comparisons showed that when the outcome was positive (+5%), participant felt the best if the project was Green and the worst when the project was Red (Green_gain: 1.57 ± 0.15, Neutral_gain: 1.28 ± 0.11, Red_gain: 1.01 ± 0.15; Green_gain vs. Neutral_gain: *p* = 0.021; Red_gain vs. Neutral_gain: *p* = 0.038; Green_gain vs. Red_gain: *p* = 0.002). When the outcome was negative (−5%), participants felt the best if the project was Red (Green_loss: −1.06 ± 0.18, Neutral_ loss: −0.97 ± 0.12, Red_loss: −0.74 ± 0.16; Green_ loss vs. Neutral_ loss: *p* = 0.458; Red_ loss vs. Neutral_loss: *p* = 0.115; Green_loss vs. Red_loss: *p* = 0.014) (Fig. 2B). Additionally, to explore whether the pleasantness rating during the outcome phase was correlated with the behavioral bias during the decision stage, we ran a Pearson’s correlational analysis between the behavioral index and the morality effect on outcome ratings (*Pleasantness Rating for Green _Gain>Loss_ > Red _Gain>Loss_*). Results showed that they were significantly correlated with each other (*r* = 0.659, *p* < 0.001).

### fMRI results

#### Decision Phase

##### Brain Activity

The contrast between trials in which the participant invested ¥1000 or ¥100 revealed significant activation in five clusters, with their peaks at left anterior cingulate gyrus (MNI coordinate [−9, 39, 21]), right cerebellum ([12, −69, −36]), left thalamus ([0, −12, 0]), left orbital part of the inferior frontal gyrus ([−27, 24, −15]), and right fusiform gyrus ([24, −87, −9]) (Table 1).

**Table 1.** Whole-brain results for choosing ¥1,000 > choosing ¥100 during the decision phase.

| Region | Brodmann<br>Area | MNI Coordinates |  |  | No. of<br>Voxels | t-value |
| --- | --- | --- | --- | --- | --- | --- |
|  |  | x | y | z |  |  |
| Left Anterior Cingulate Gyrus | 32 | -9 | 39 | 21 | 97 | 6.06 |
| Right Anterior Cingulate Gyrus | 32 | 9 | 36 | 21 |  | 3.97 |
| Left Anterior Cingulate Gyrus | 32 | -3 | 42 | 6 |  | 3.96 |
| Right Cerebellum |  | 12 | -69 | -36 | 207 | 5.31 |
| Right Cerebellum |  | -24 | -60 | -48 |  | 5.10 |
| Right Cerebellum |  | -18 | -45 | -45 |  | 4.88 |
| Left Thalamus |  | 0 | -12 | 0 | 160 | 5.10 |
| Brain Stem |  | 6 | -27 | -12 |  | 4.67 |
| Right Thalamus |  | 6 | -18 | 3 |  | 4.50 |
| Left Inferior Frontal Gyrus (orbital part) | 47 | -27 | 24 | -15 | 63 | 4.48 |
| Left Insula | 48 | -33 | 15 | -12 |  | 4.35 |
| Left Superior Temporal Pole |  | -36 | 15 | -21 |  | 3.86 |
| Right Fusiform Gyrus | 18 | 24 | -87 | -9 | 62 | 4.37 |
| Right Fusiform Gyrus | 17 | 12 | -93 | 3 |  | 4.23 |
| Right Calcarine Fissure | 18 | 18 | -96 | -3 |  | 4.19 |

We considered these significantly activated clusters as ROIs and used them to determine which brain regions were sensitive to moral valence among brain circuits processing investment preferences. Within the spheres centered at these five peaks, we calculated mean parameter-estimates of the three different moral valences of the investment categories in GLM2. One-way ANOVA revealed that only the right fusiform gyrus showed gradually increasing activation strength from Red (3.54 ± 0.72) to Neutral (4.08 ± 0.68) to Green investments (4.50 ± 0.73) (*F*_(2,32)_ = 10.04, *p* < 0.001; Fig. 3A). Post-hoc pairwise t-tests with Bonferroni correction revealed that brain activity for Neutral investments was significantly higher than that for Red investments (*t*_(32)_ = 2.55, *p* = 0.016) and significantly lower than that for Green investments (*t*_(32)_ = 2.47, *p* = 0.019) (Fig. 3A).

**Figure 3.**
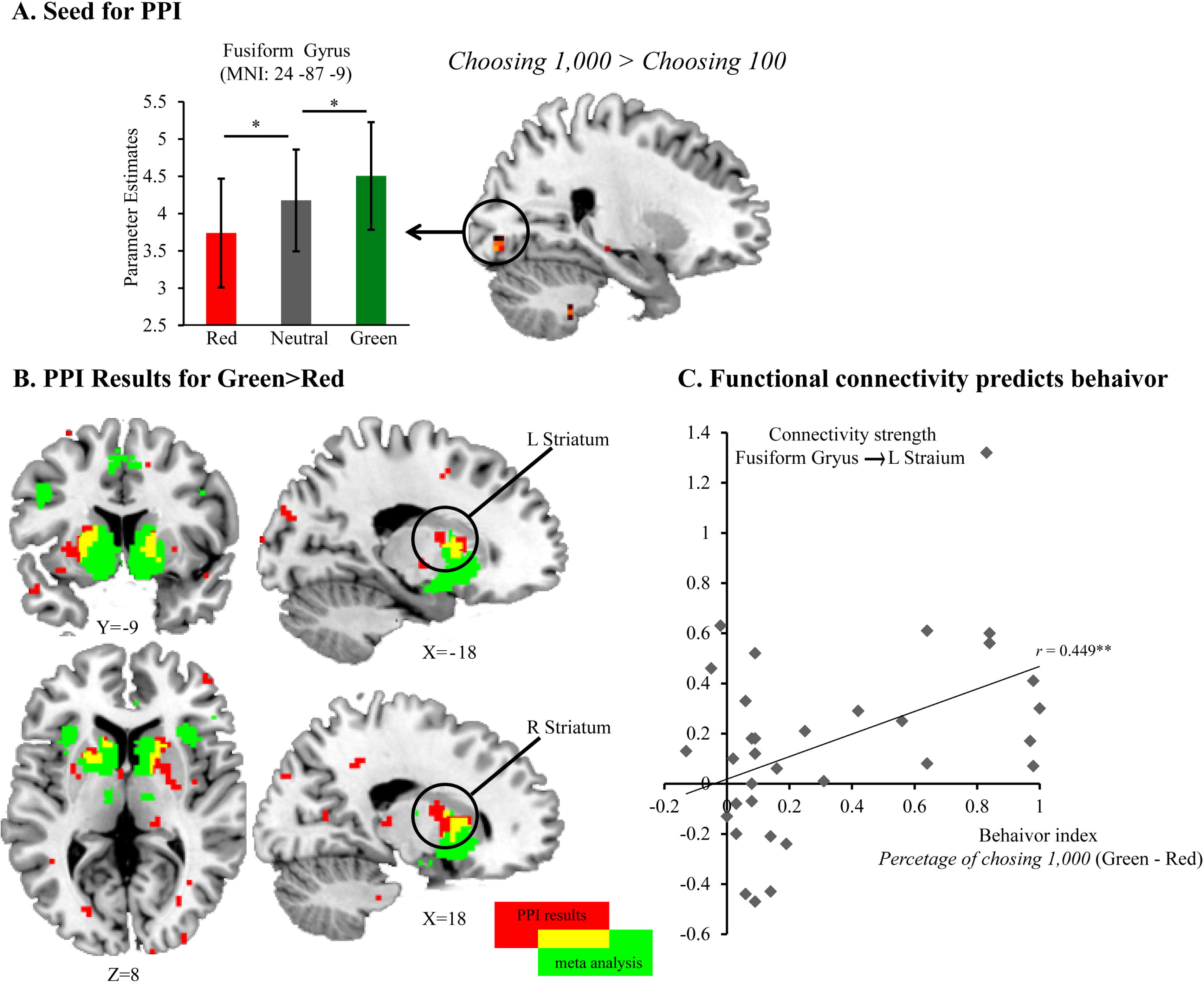
fMRI results during the decision phase. **(A)** The fusiform gyrus was revealed in the Choosing 1,000 > Choosing 100 contrast and was sensitive to moral valence of the investment category. **(B)** PPI analysis with the fusiform gyrus as the seed region on the contrast of Green>Red revealed bilateral striatum. Overlap with reward circuitry as defined in a meta-analysis is also shown (red: PPI results; green: reward circuitry in the meta-analysis; yellow: the overlap). **(C)** The strength of the functional connectivity between the fusiform gyrus and left striatum predicted the behavioral index.

##### Functional Connectivity

In the previous activation analysis, we found that among brain circuits processing investment preferences, the right fusiform gyrus was sensitive to moral valence. We considered the right fusiform gyrus ([24, −87, −9]) as a seed region and calculated the functional connectivity contrast between Green and Red investments using PPI. The results revealed significantly stronger functional connectivity between the right fusiform gyrus and the left putamen ([24, −87, −9]), right nucleus accumbens ([18, 0, 12]), and left cuneus ([−9, −75, 24]) for Green investments than for Red investments (Table 2).

**Table 2.**
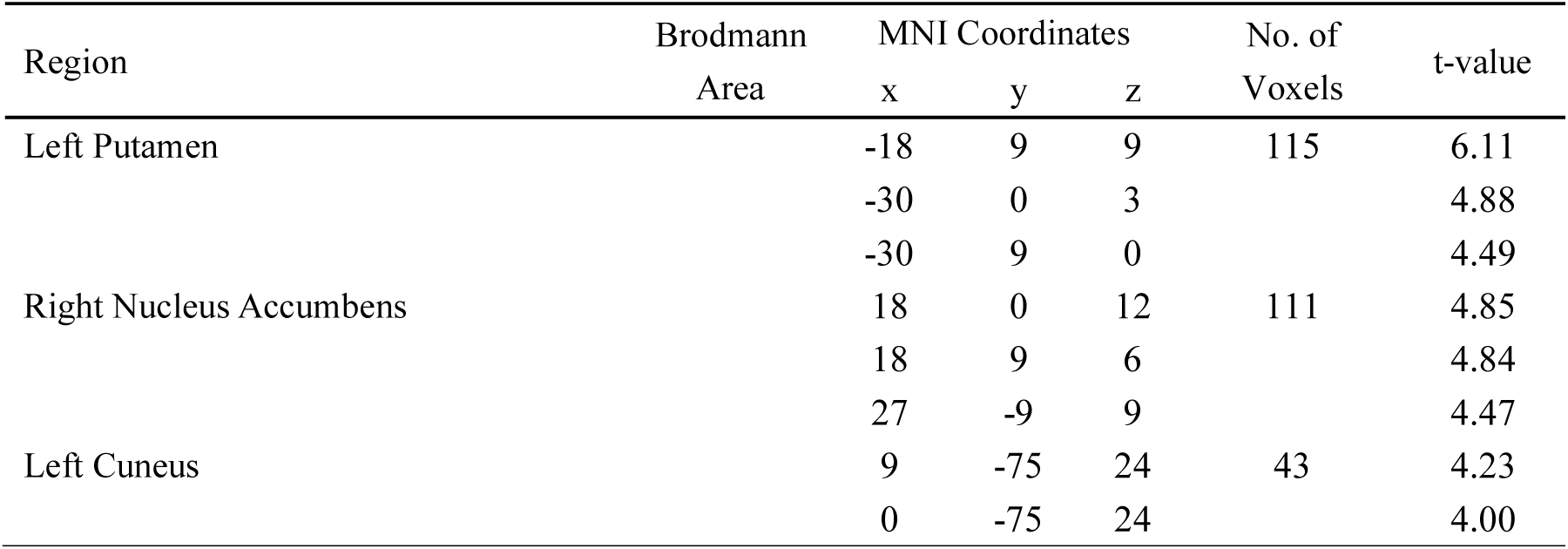
PPI Results. Loci showing significantly different functional connectivity based on the Green> Red contrast of investment category during the decision phase

| Region | Brodmann<br>Area | MNI Coordinates |  |  | No. of<br>Voxels | t-value |
| --- | --- | --- | --- | --- | --- | --- |
|  |  | x | y | z |  |  |
| Left Putamen |  | -18 | 9 | 9 | 115 | 6.11 |
|  |  | -30 | 0 | 3 |  | 4.88 |
|  |  | -30 | 9 | 0 |  | 4.49 |
| Right Nucleus Accumbens |  | 18 | 0 | 12 | 111 | 4.85 |
|  |  | 18 | 9 | 6 |  | 4.84 |
|  |  | 27 | -9 | 9 |  | 4.47 |
| Left Cuneus |  | 9 | -75 | 24 | 43 | 4.23 |
|  |  | 0 | -75 | 24 |  | 4.00 |

We downloaded the results of a rewards-processing meta-analysis from Neurosyth and considered it as a mask. Two clusters of significant PPI results overlapped with the mask, namely the left putamen ([−18, 6, 9]) and the right nucleus accumbens ([18, 9, 6]) (Fig. 3B). We extracted the mean parameters of the functional connectivity maps within spheres centered at these peaks and found that the functional connectivity between right fusiform gyrus and left putamen could significantly predict the behavioral index (*r* = −0.449, *p* = 0.009) after Bonferroni correction. The functional connectivity between right fusiform gyrus and right nucleus accumbens predicted the behavioral index in the outcome phase, although significance was marginal (*r* = 0.343, *p* = 0.051) and did not survive Bonferroni Correction (Fig. 3C).

#### Outcome phase

Two types of second-level analyses were considered in the outcome phase. We first performed one-sample t-tests to explore whether brain activity was different for Red and Green investments. The results showed no significant differences between the two conditions. Then, we considered the behavioral index as a regressor and regressed it against the Gain > Loss contrast at the group level. The results indicated significant activation in the left cerebellum ([−6, −57, −15]), brain-stem ([−3, −27, −12]), right orbital part of the middle frontal gyrus ([39, 30, −3]), right superior medial frontal gyrus ([3, 42, 45]), right middle cingulate gyrus ([3, 30, 33]), and right putamen ([18, 12, 0]) (Table 3 and Fig. 4A).

**Figure 4.**
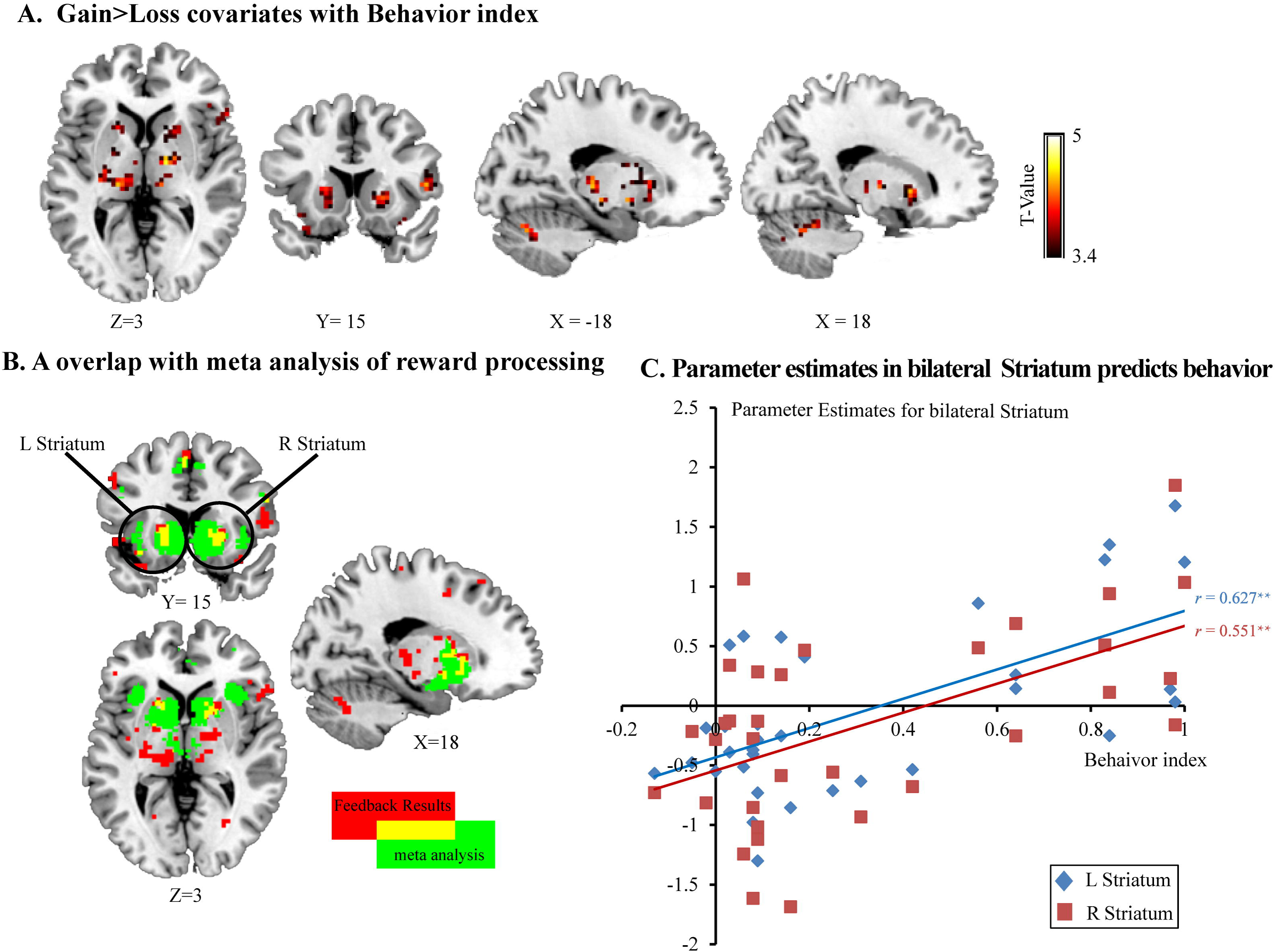
fMRI results during the outcome phase. **(A)** Activity levels in regions that passed the Gain>Loss contrast threshold correlated positively with the behavioral index at the group level. **(B)** The regions that covaried with individual behavioral indexes overlapped with the reward circuitry that was revealed in the met-analysis (red: results of **(A)**; green: reward circuitry in the meta-analysis; yellow: the overlap). **(C)** Mean parameter estimates for bilateral striatum predicted the behavioral index.

**Table 3.** Regions revealed in the Gain>Loss contrast covaried with the individual behavioral index during the outcome phase

| Region | Brodmann<br>Area | MNI Coordinates |  |  | No. of<br>Voxels | t-value |
| --- | --- | --- | --- | --- | --- | --- |
|  |  | x | y | z |  |  |
| Left Cerebellum | 18 | -6 | -57 | -15 | 289 | 6.12 |
| Right Cerebellum |  | 12 | -57 | -24 |  | 6.02 |
| Right Cerebellum | 18 | 12 | -66 | -18 |  | 4.70 |
| Brain Stem |  | -3 | -27 | -12 | 436 | 5.82 |
| Right Thalamus |  | 12 | -24 | 9 |  | 5.01 |
| Right Middle Frontal Gyrus (orbital) | 47 | 39 | 30 | -3 | 193 | 5.71 |
| Right Insula | 47 | 30 | 21 | -12 |  | 5.04 |
| Right Inferior Frontal Gyrus (Triangular) | 45 | 57 | 24 | 6 |  | 5.02 |
| Right Superior Medial Frontal Gyrus | 9 | 3 | 42 | 45 | 46 | 4.91 |
| Right Superior Medial Frontal Gyrus | 9 | 12 | 36 | 48 |  | 4.84 |
| Right Middle Cingulate Gyrus | 24 | 3 | 30 | 33 | 186 | 4.84 |
| Right Anterior Cingulate Gyrus | 32 | 6 | 42 | 27 |  | 4.75 |
| Right Middle Cingulate Gyrus | 32 | 12 | 27 | 33 |  | 4.61 |
| Right Putamen |  | 18 | 12 | 0 | 73 | 4.44 |
| Right Amygdala | 34 | 24 | 0 | -12 |  | 3.90 |
| Right Putamen | 34 | 30 | -6 | -9 |  | 3.84 |

We then masked that result with the meta-analysis result for reward from Neurosyth and localized several ROIs among the mask including left putamen ([−18, 15, 3]), right putamen ([18, 12, 0]), right orbital part of the inferior frontal gyrus ([39, 30, −3]), right middle cingulate gyrus ([3, 30, 33]), left insula ([−30, 18, −9]), and left thalamus ([−9, −21, 9] and [−6, −15, −9]) (Fig. 4B). We extracted the mean parameters of the brain activation maps within spheres centered at these peaks and found that brain activity in the left and right putamen could significantly predict the behavioral index during the outcome phase (left: *r* = 0.551, *p* = 0.001; right: *r* = 0.627, *p* < 0.001) after Bonferroni correction (Fig. 4C).

## Discussion

This study shows that morality modulates economic decision making and that brain activity and functional connectivity within reward circuitry are likely related to this modulating effect, both when making decisions and when observing the outcomes.

Behaviorally, in a profit-seeking hypothetical investment game, participants were significantly influenced by the moral valence of investment category such that they invested the most in the moral categories (*Green*) and the least in the immoral ones (*Red*), even though the chance of gaining money was the same. The moral valence of investments also influenced how participants felt about the outcomes such that having chosen Green investments triggered the highest feeling of pleasantness for gains and the lowest for losses. In contrast, having chosen Red investments triggered the lowest feelings of pleasantness for gains and the highest for losses. Moreover, the modulating effect of morality measured during the decision phase was significantly correlated with its effect measured during the outcome phase, suggesting a consistent, prevalent influence of morality.

BOLD signals acquired during the decision phase revealed that activation of the higher-level visual processing cortex, specifically the fusiform gyrus (FG), was not only correlated with behavioral choice but was also sensitive to the moral valence of the project such that Green projects triggered the strongest activation while Red projects triggered the weakest. This finding is interesting because it suggests that moral valence can be encoded during visual processing, which is compatible with the previously reported involvement of visual encoding in moral reasoning and judgments (Amit and Greene, 2012; Garon et al., 2018), and in the moral emotions such as guilt and shame (Michl et al., 2014). An event-related potential (ERP) study combined with source estimations revealed stronger activation of the FG when visually presented adjectives were sent by a human than when they were sent by a computer as well as when the words used were emotional than when they were emotionless. The FG is thus suggested as a primary generator of social and emotional effects during visual word processing (Schindler et al., 2015). Enhanced FG activity has been suggested to reflect enhanced attention and visual imagery (Boccia et al., 2017), and therefore can facilitate the social motivation of the stimuli (Utevsky et al., 2017). In the present study, relative to the Neutral category, the higher activation for Green investments and the lower activation for Red investments might reflect modulated attention allocation and social motivations by morality during the visual processing. Green investments might draw more attention and trigger more active visual imagery (e.g., visualizing the benefits to society or positive social rewards that they might gain through the investment), which would imbue with greater motivational relevance. In contrast, Red investments might trigger avoidance tendencies and result in the opposite effect.

Analysis of functional connectivity further revealed that for Green investments, connectivity between the FG and reward circuitry, including the left putamen and right nucleus accumbens, was significantly stronger than for Red investments. Moreover, the magnitude of the connectivity between the FG and left putamen was able to predict the behavioral tendency (i.e., investing more in the Green/moral direction than in the Red/immoral direction). The striatum has been consistently found to be activated during both expecting and experiencing monetary (Bartra et al., 2013) and social rewards, with higher rewards inducing greater activity (Izuma et al., 2008; Rademacher et al., 2010; Schultz et al., 2019; Spreckelmeyer et al., 2009). In the present study, Green investments contained a positive moral value such as benefitting others and society and bringing positive social judgments for the investor. Red investments inherited negative moral values. Thus, the enhanced connectivity between the FG and ventral striatum (the right nucleus accumbens) may reflect enhanced subjective values for Green investments and decreased subjective values for Red investments due to the presence of moral/immoral values.

Additionally, recent studies have highlighted the role of the dorsal striatum (including putamen) in the learning of action–reward associations from feedback (Balleine et al., 2007; Delgado, 2007; Goulet-Kennedy et al., 2016). As we discuss in the following paragraph, morality also modulates behavioral and neural responses to the processing of decision outcomes. Thus, feedback learning might also contribute to the difference in connectivity strength between Green and Red directions.

BOLD signals acquired during the outcome phase revealed that the differences in neural responses between gain and loss within the reward network (bilateral putamen, left caudate, right inferior gyrus, bilateral insula, left medial cingulate cortex, and left thalamus) covaried with the behavioral index, indicating the strength of the morality effect on a between-participant level. Thus, greater differences in BOLD activation between gain and loss within the reward network is associated with greater behavioral bias caused by morality (i.e., investing more in Green projects than in Red ones). These results were consistent with subjective ratings of pleasantness such that Green investments enlarged the discrepancy between gain and loss while Red investments showed the opposite effect. When investing in Green projects, participants not only expected economic profits but also a positive moral value. In contrast, when investing in Red projects, a negative moral value would be generated along with the expectations of economic profit. This difference in moral value may underlie the observed effect within the reward circuitry as well as the levels of pleasantness that were reported.

Previous studies have suggested that activation in the striatum when experiencing rewards can predict decision behavior, probably through feedback-learning. For instance, smokers who displayed the weakest activity in the ventral striatum when receiving monetary rewards were less keen to refrain from smoking for monetary reinforcement (Wilson et al., 2014). Although the present experimental design could not directly test whether morality modulates feedback-learning, two pieces of evidence support this hypothesis. First, during the outcome phase, morality modulates the difference between gain and loss in both the brain (BOLD activation) and behavior (pleasantness rating) in the same manner (i.e., Green enlarged the Gain>Loss difference while Red reduced it). The morality effect measured during the outcome phase was significantly correlated with that measured during the decision phase, indicating the existence of the feedback learning loop and the consistent modulating effect of morality. Second, we found that participants gained the most when they invested in Green categories and the least when they invested in Red categories. This is consistent with the findings that Green categories enlarged the difference between gain and loss and thus facilitated the learning process, while the Red categories showed the opposite effect.

The decision to invest larger sums of money can be considered a preference for higher risk because investing more money increases the potential gains and losses. On the flip side, investing smaller sums indicates a preference for lower risk. In this sense, the present results can be explained as morality modulating the tendency to take risks such that Green projects made participants more prone to take risks than did Red projects. However, because we found that participants made significantly more profit in Green investments than in Red investments, we suggest that the positive moral valence facilitates the probability learning thus resulted in higher gains while the negative moral valence harmer it and resulted in lower gains would be a more possible explanation, comparing to the modulating of risk preference.

Our study provides evidence that morality modulates both the decision making and the outcome evaluation in economic situations; people are willing to invest a larger amount of money into a moral project that may benefit society than they are into an immoral project that they think will harm society. They also rate gains in moral investments as more pleasant and losses as the most unpleasant. In the brain, we found that the reward system, especially the bilateral striatum, was involved in modulating functional connectivity during both phases, but in different ways. During decision making, the functional connectivity between fusiform gyrus and striatum might underlie the observed investing bias (Green over Red projects), while the covariation of BOLD signals in bilateral striatum with the behavioral tendency might explain the effect observed during the outcome evaluations.

One limitation of the present study is that the ecological validity of the design was not high because the hypothetical investment task did not actually resemble real investment. Outcomes for most investments are delayed rather than immediate. In reality, many Red projects can make “fast” money while Green projects may take longer to make a profit or show their benefits. However, we did not take differences in monetary gain delays between projects into consideration. Another limitation is that the experimental design did not have enough trials to explore how morality modulates the feedback learning process, which would be an interesting and meaningful question for further studies.

## Author Contributions

Author contributions: F.C. designed the research; C.L. and X.H collected the data; J.L analyzed data; F.C., and J.L wrote the paper. The authors declare no competing financial interests.

## Acknowledgment

This work was by the National Natural Science Foundation of China (no. 31871109 to F.C. and no. 31900779 to J.L).

